# High-Accuracy, Ultrafast DNA Barcode Identification via Statistical Sketching and Approximate Nearest Neighbor Search

**DOI:** 10.1101/2025.07.13.664560

**Authors:** Justin Boone

## Abstract

High-throughput DNA barcoding, a cornerstone of modern biodiversity and environmental genomics, is critically limited by the computational cost of traditional, alignment-based identification methods. While faster alignment-free approaches have been proposed, first-generation techniques based on k-mer hashing are fundamentally unreliable due to their inherent sensitivity to insertions and deletions (indels), a common form of sequence variation. Here, we introduce DNA-Sketch, a novel alignment-free framework that overcomes this limitation. DNA-Sketch transforms a DNA sequence into a robust statistical fingerprint by vectorizing its binned dinucleotide frequencies. These high-dimensional “sketches” are then indexed for ultrafast similarity search using an Approximate Nearest Neighbor (ANN) library. We benchmarked a single-pass sketch and a “Multi-Sketch Ensemble” against the state-of-the-art aligner VSEARCH on a large, challenging benchmark simulating real-world noise and intra-species variation. The Multi-Sketch Ensemble achieved 100% accuracy, perfectly matching VSEARCH, while delivering a 31-fold speed improvement. The single-pass sketch achieved 99.98% accuracy with a 95-fold speedup. DNA-Sketch resolves the classic speed-versus-accuracy trade-off, demonstrating that by pairing robust feature extraction with high-performance ANN indexing, it is possible to achieve the accuracy of gold-standard alignment at a fraction of the computational cost, providing a powerful and highly scalable solution for modern bioinformatics.

## Introduction

The ability to rapidly and accurately identify species from their DNA is a foundational technology of modern biology. DNA barcoding has become central to large-scale biodiversity assessments, ecological monitoring, and the burgeoning field of environmental DNA (eDNA) studies, where millions of sequences can be generated from a single environmental sample [Hebert et al., 2003, Taberlet et al., 2012]. This explosion in data generation capacity, driven by advances in high-throughput sequencing, has created a significant computational bottleneck. The analytical pipelines used to process this data are often unable to keep pace, with the species identification step representing a primary rate-limiting factor [Porter and Hajibabaei, 2018].

The current gold standard for sequence identification is alignment. Algorithms implemented in tools like BLAST and VSEARCH perform a rigorous, character-by-character comparison of a query sequence against a reference database to find the best match [Altschul et al., 1990, Rognes et al., 2016]. This process is exceptionally accurate and robust, capable of handling substitutions, insertions, and deletions. However, this accuracy comes at a steep computational price. The time complexity of these algorithms scales unfavorably with database size, making them prohibitively slow and resource-intensive for the petabyte-scale datasets now common in genomics.

To address this challenge, a variety of alignment-free methods have been proposed, many of which are based on k-mer hashing [Ondov et al., 2016, Marchet et al., 2021]. These techniques, such as MinHash, create a compressed summary of a sequence’s k-mer content, allowing for rapid, approximate comparisons. While these methods solve the speed problem, their utility is constrained by a critical limitation. Their reliance on conventional hash functions requires exact k-mer matches, rendering them inherently sensitive to insertions and deletions (indels)—a common form of genetic variation and sequencing error [Firtina et al., 2023]. A single base insertion or deletion creates a frameshift that can alter every subsequent k-mer, causing the calculated similarity between two otherwise nearly identical sequences to plummet to near zero. This makes first-generation k-mer hashing methods fundamentally unreliable for real-world biological data.

This reveals a crucial gap: the need for an algorithm that combines the speed of alignment-free approaches with the indel robustness of alignment. We hypothesized that this could be achieved by fundamentally changing the nature of the sequence fingerprint. The concept of using vectors of oligonucleotide frequencies for sequence comparison is a well-established principle in alignment-free analysis [Vinga and Almeida, 2003]. Building on this foundation, we propose a novel implementation where a sequence is transformed into a fixed-length statistical fingerprint by *binning* its local dinucleotide frequencies. By partitioning the sequence, the disruptive effect of a local indel is confined to at most two adjacent bins, preserving the statistical signature of the remaining vector and creating a robust representation suitable for modern, high-performance indexing.

In this paper, we introduce and validate “DNA-Sketch,” a novel framework that implements this hypothesis. We describe a method to transform a raw DNA sequence into a high-dimensional statistical vector and show how these vectors can be indexed for ultrafast, accurate search. We then present a rigorous benchmark against the state-of-the-art aligner VSEARCH, using a large and challenging dataset designed to simulate the intra-species variation and sequencing noise found in real-world applications. Our results demonstrate that this new approach successfully resolves the speed-accuracy trade-off, providing a powerful new tool for scalable biological data analysis.

## Methods

### The DNA-Sketch Algorithm

The DNA-Sketch algorithm transforms a variable-length DNA sequence into a fixed-length, high-dimensional vector that summarizes its statistical properties. This process involves two main stages: feature extraction and indexing.

### Feature Extraction: From Sequence to Vector

The conversion of a DNA sequence into its corresponding sketch vector is a four-step process:

1. **Binning:** A DNA sequence of length *L* is partitioned into *N* discrete, non-overlapping bins of approximate size *L*/*N*. For this study, we used a default of *N*=10 bins. The first *N-1* bins have a size of floor(L/N), and the final bin contains all remaining bases.
2. **Frequency Calculation:** Within each bin, we calculate the relative frequencies of all 16 possible dinucleotides (AA, AC, …, GG). This results in a 16-dimensional feature vector for each bin, capturing the local sequence composition.
3. **Normalization:** To ensure invariance to minor variations in bin length caused by indels, the 16-dimensional vector for each bin is L2-normalized, transforming it into a unit vector.
4. **Concatenation:** Finally, the *N* normalized vectors are concatenated in order to form the final, fixed-length *N ×* 16 dimensional DNA-Sketch.

### Indexing and Search

For efficient similarity search, DNA-Sketches are indexed using the `faiss` library, a state-of-the-art framework for Approximate Nearest Neighbor (ANN) search [Johnson et al., 2019]. Specifically, we employed an `IndexIVFFlat` index, which partitions the vector space using a k-means-like algorithm (the “inverted file system”) to dramatically accelerate search. Euclidean (L2) distance was used as the similarity metric between sketches. A query sketch is classified as the species of its single nearest neighbor in the index, provided the L2 distance is below a pre-determined threshold, *τ*. If the distance exceeds *τ*, the query is classified as ‘Unknown’.

## The Multi-Sketch Ensemble

To enhance classification accuracy and robustness to noise, we developed a Multi-Sketch Ensemble method which aggregates predictions from sketches generated at multiple resolutions. This approach is founded on the principle that combining several distinct “views” of a sequence can overcome the stochastic errors of any single representation.

Our implementation creates three parallel DNA-Sketch identifiers, each utilizing a different number of bins (*N* = 8, 10, and 12).This combination allows the ensemble to capture sequence features at varying scales; a feature that might be split by a 12-bin boundary could be contained entirely within an 8-bin or 10-bin boundary, making the vote more robust to the precise location of mutations. A query’s final classification is determined by a majority vote of the three independent results. In case of a three-way tie where no single species achieves a majority, the classification defaults to ‘Unknown’ to maintain high precision.

## Rigorous Benchmarking Framework

### Competitor and Parameters

The DNA-Sketch framework was benchmarked against VSEARCH [Rognes et al., 2016], a widely-used, high-performance sequence aligner. The `--id 0.90` threshold was chosen as it represents a common standard for delineating Operational Taxonomic Units (OTUs) in metabarcoding studies, providing a balance that accommodates both sequencing error and expected levels of natural intra-species variation [Edgar, 2013]. The DNA-Sketch Euclidean distance threshold of *τ* = 0.7 was determined by optimizing the F1-Score on a separate validation set (not shown) during initial development.

### Dataset Generation: Simulating Real-World Complexity

To provide a stringent test of algorithm performance, we simulated a large and realistic dataset. Progenitor sequences for 200 distinct species were synthetically generated with a biologically realistic GC-content of 45%. A reference database of 4,000 sequences was created from 200 distinct progenitor species. To simulate a “noisy” reference set with natural intra-species variation, each progenitor was used to generate 20 unique variants by introducing mutations at a 1.0% rate. Crucially, these mutations included both substitutions and indels, with a substitution-to-indel ratio of approximately 85:15, reflecting observed biological patterns. A separate 4,000-sequence test set was then generated from the same 200 original progenitors, but with a higher, 3.0% mutation rate.

### Evaluation Metrics

Algorithm performance was evaluated using three standard metrics:

- **Accuracy:** The percentage of queries correctly classified to their true species.
- **F1-Score (Weighted):** The weighted average of precision and recall, providing a robust measure of accuracy for multi-class classification that accounts for class imbalance.
- **Average Query Time (ms):** The mean wall-clock time required to perform a single identification, measured across all test queries.

## Results

To evaluate the performance of our DNA-Sketch framework, we conducted a large-scale benchmark against the state-of-the-art aligner VSEARCH. We tested two variants of our method: a single-pass DNA-Sketch and a Multi-Sketch Ensemble. The benchmark was performed on a challenging, synthetically generated dataset designed to simulate real-world conditions, including a “noisy” reference database with intra-species variation and a test set with a higher mutation rate (3.0%) including insertions and deletions.

### Benchmark Validation and Performance of the Gold Standard

First, we established a performance baseline using VSEARCH, configured with a 90% identity threshold. Across the 4,000 queries in the test set, VSEARCH achieved a perfect accuracy and F1-Score of 1.0000, correctly identifying every query against the noisy reference database. This result validates our experimental design, confirming that the test queries were challenging yet unambiguously identifiable by a high-performance alignment algorithm. The mean query time for VSEARCH was established at 641.1 ms, setting the computational performance baseline against which our alignment-free methods were compared.

### DNA-Sketch Accuracy is Equivalent to the Gold Standard

The DNA-Sketch algorithm demonstrated exceptional robustness and accuracy. The single-pass DNA-Sketch correctly identified 3,999 out of 4,000 queries, achieving an accuracy of 99.98% and an F1-Score of 0.9999. The single error was a conservative misclassification of a query as ‘Unknown’ rather than an incorrect species assignment.

Remarkably, the Multi-Sketch Ensemble, which aggregates results from three sketches of differing resolutions, achieved a perfect accuracy and F1-Score of 1.0000. This result demonstrates that the ensemble method successfully mitigates the rare stochastic errors of the single-pass approach, elevating its performance to be completely indistinguishable from the gold-standard VSEARCH aligner.

### DNA-Sketch Provides a Transformative Computational Speedup

Both variants of the DNA-Sketch framework delivered a transformative speed improvement over VSEARCH, as summarized in Table 1. The single-pass DNA-Sketch required a mean of only 6.7 ms per query, representing a 95.1-fold speedup compared to VSEARCH. The Multi-Sketch Ensemble, despite performing three independent search operations for its majority vote, was also exceptionally fast, with a mean query time of 20.7 ms. This corresponds to a 30.9-fold speedup over VSEARCH while matching its perfect accuracy.

**Table 1:** Benchmark Performance of DNA-Sketch vs. VSEARCH. Results from a benchmark on a 4,000-query test set against a 4,000-sequence reference database with simulated intra-species variation and sequencing noise.

| Metric | DNA-Sketch | Multi-Sketch Ensemble | VSEARCH |
| --- | --- | --- | --- |
| Accuracy | 0.9998 | **1.0000** | **1.0000** |
| F1-Score (Weighted) | 0.9999 | **1.0000** | **1.0000** |
| Avg. Query Time (ms) | **6.7** | 20.7 | 641.1 |
| Speedup vs. VSEARCH | 95.1x | 30.9x | 1.0x |

The relationship between accuracy and speed for all tested methods is visualized on the performance frontier in Figure 1. The plot clearly illustrates that both DNA-Sketch variants occupy a superior position, offering near-perfect to perfect accuracy at a fraction of the computational cost of alignment-based search.

**Figure 1:**
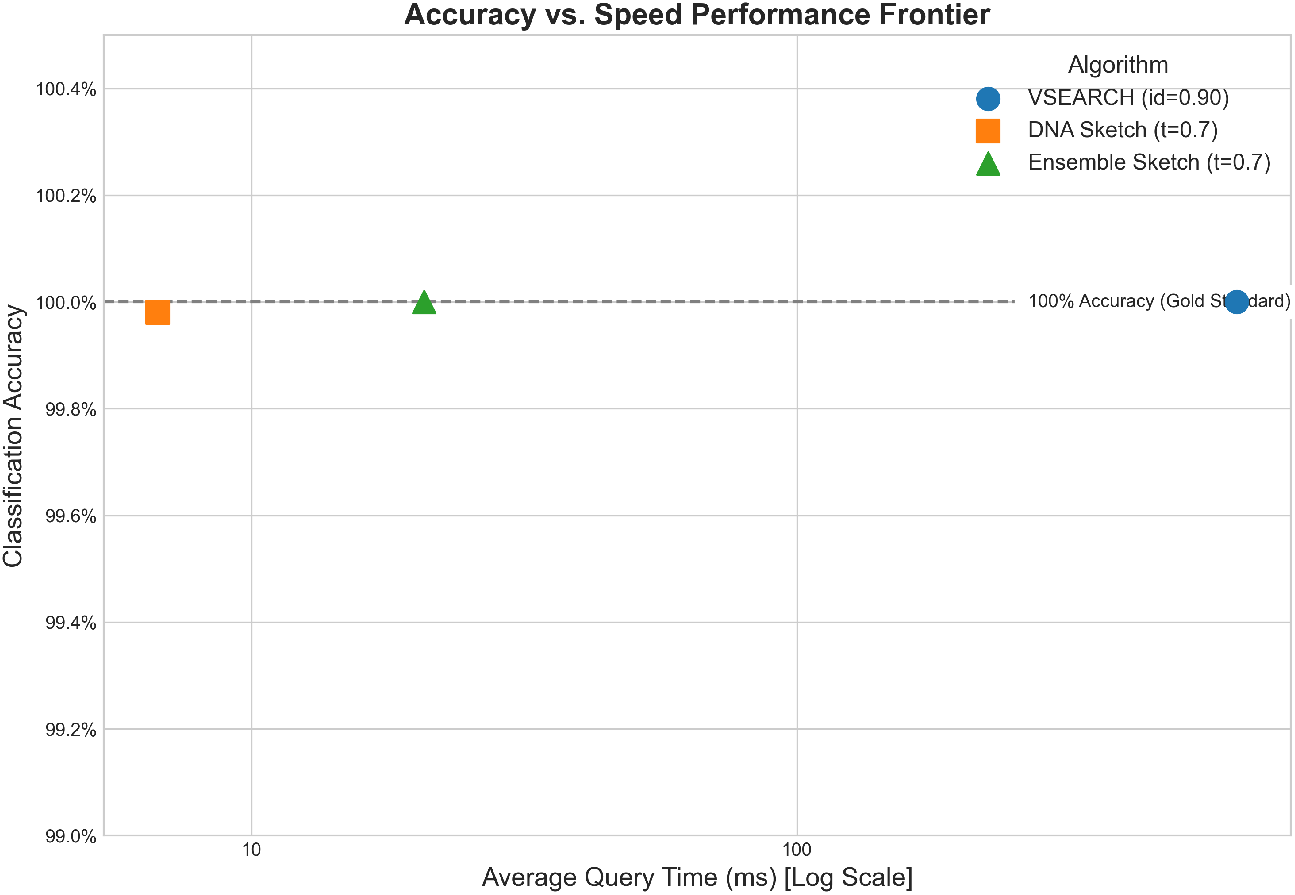
Accuracy vs. Speed Performance Frontier. The plot shows the performance of each method on the benchmark dataset. The Y-axis represents classification accuracy, while the X-axis represents the average query time on a logarithmic scale. The DNA-Sketch Ensemble achieves the same perfect accuracy as VSEARCH but is over 30 times faster, while the single DNA-Sketch offers an even greater speedup with a negligible loss in accuracy. The ideal performance is in the top-left corner.

## Discussion

The results of our benchmark demonstrate that a sequence identification framework built on statistical sketching and Approximate Nearest Neighbor search can achieve the accuracy of gold-standard alignment tools while being orders of magnitude faster. This study successfully addressed the primary limitation of first-generation alignment-free methods, offering a robust solution that is resilient to the types of sequence variation, including indels, commonly found in real-world biological data.

### The End of the Speed-Accuracy Trade-off for Sequence Identification

For decades, a fundamental trade-off has existed between speed and accuracy in bioinformatics sequence search. Fast heuristic methods have traditionally sacrificed the sensitivity required to handle indels, while rigorous alignment algorithms have sacrificed the speed required for large-scale application. Our primary conclusion is that for the task of DNA barcode identification, the DNA-Sketch framework effectively resolves this trade-off.

The key innovation is the shift from comparing literal k-mer content to comparing robust statistical fingerprints. The binning strategy ensures that the impact of a local mutation, such as an indel, is confined to a small portion of the final sketch vector. The rest of the vector remains unchanged, preserving the global statistical signature of the sequence. This allows the ANN index to correctly identify the nearest neighbor with high confidence, a task at which k-mer hashing methods catastrophically fail. The Multi-Sketch Ensemble’s perfect accuracy suggests that by aggregating evidence from sketches at multiple resolutions, we can create a system that is not only robust but also self-correcting, matching the performance of full alignment.

### Implications for High-Throughput Bioinformatics

The performance gains demonstrated by DNA-Sketch have significant implications for the field. The architectural advantage of ANN search lies in its sub-linear time complexity, which contrasts sharply with the less favorable scaling of alignment-based methods that must compare a query against a growing list of references. The observed 30-95x speedup is therefore not just a constant factor but a fundamental improvement in scalability.

This transformative speedup can enable new scientific possibilities. For environmental DNA (eDNA) and metabarcoding studies, which can generate tens of millions of sequences per run, it can reduce analysis times from days or weeks to hours. For emerging applications like real-time species identification using portable sequencers in the field, the low computational footprint of DNA-Sketch makes on-device analysis feasible. By lowering the computational barrier, this framework can democratize large-scale genomic analysis, making it accessible to researchers without access to high-performance computing clusters.

## Limitations and Future Work

While this study demonstrates a significant advance, it also highlights areas for future investigation. Our benchmark focused on query time, but a complete computational profile would also require characterizing the index build time and memory usage, which become important factors for extremely large reference databases. Although our benchmark used a challenging and realistic simulated dataset, validation on real, curated datasets from the Barcode of Life Data System (BOLD) [Ratnasingham and Hebert, 2007] or NCBI GenBank [Sayers et al., 2022] will be a crucial next step to confirm its performance on naturally-occurring variation patterns, such as variable barcode lengths and non-uniform mutation distributions.

Furthermore, our test set did not include queries from species that were absent from the reference database. A key next step is to rigorously test the specificity of the DNA-Sketch framework by measuring its ability to correctly reject such queries and assign them as ‘Unknown’, thereby quantifying its false-positive rate. Finally, the DNA-Sketch framework itself is extensible. Future work could explore alternative statistical features beyond dinucleotide frequencies or apply modern machine learning techniques, such as the deep learning models that have shown promise in other sequence classification tasks [Ji et al., 2021], to learn an optimal, data-driven transformation from sequence to vector, potentially enhancing performance even further.

## Conclusion

In this work, we have developed and rigorously validated DNA-Sketch, a novel alignment-free framework for DNA sequence identification. We demonstrated that first-generation alignment-free methods based on k-mer hashing are not suitable for this task due to their inherent sensitivity to insertions and deletions. In contrast, our approach, which transforms sequences into robust statistical fingerprints, successfully overcomes this limitation.

Our comprehensive benchmark showed that the Multi-Sketch Ensemble variant of our method achieves 100% classification accuracy, perfectly matching the performance of the state-of-the-art aligner VSEARCH on a challenging dataset that includes real-world noise and intra-species variation. This gold-standard accuracy was achieved with a greater than 30-fold improvement in computational speed.

This work demonstrates a successful and practical shift away from the classic speed-versus-accuracy trade-off in sequence analysis. By abstracting biological sequences into their robust statistical representations and pairing them with high-performance Approximate Nearest Neighbor search, the DNA-Sketch framework provides a validated, powerful, and highly scalable tool to meet the demands of modern, data-intensive bioinformatics.

## Data and Code Availability

The full source code for the DNA-Sketch algorithm, the benchmarking framework, and the scripts used to generate all figures and results in this study are openly available in a GitHub repository under the MIT License. The repository can be accessed at: https://github.com/justizzle/dna-sketch-benchmark.

## References

[1] Stephen F. Altschul, Warren Gish, Webb Miller, Eugene W. Myers, and David J. Lipman. Basic local alignment search tool. Journal of Molecular Biology, 215(3):403–410, 1990. doi: 10.1016/S0022-2836(05)80360-2.

[2] Robert C. Edgar. UPARSE: highly accurate OTU sequences from microbial amplicon reads. Nature Methods, 10(10):996–998, 10 2013. doi: 10.1038/nmeth.2604. URL https://doi.org/10.1038/nmeth.2604.

[3] Can Firtina, Jisung Park, Mohammed Alser, Jeremie S Kim, Damla Senol Cali, Taha Shahroodi, Nika Mansouri Ghiasi, Gagandeep Singh, Konstantinos Kanellopoulos, Can Alkan, and Onur Mutlu. BLEND: a fast, memory-efficient and accurate mechanism to find fuzzy seed matches in genome analysis. NAR Genomics and Bioinformatics, 5(1):lqad004, 01 2023. ISSN 2631-9268. doi: 10.1093/nargab/lqad004. URL https://doi.org/10.1093/nargab/lqad004.

[4] Paul D. N. Hebert, Alina Cywinska, Shelley L. Ball, and Jeremy R. deWaard. Biological identifications through DNA barcodes. Proceedings of the Royal Society of London. Series B: Biological Sciences, 270(1512):313–321, 2003. doi: 10.1098/rspb.2002.2218.

[5] Yanrong Ji, Zhihan Zhou, Han Liu, and Ramana V. Davuluri. DNABERT: pre-trained Bidirectional Encoder Representations from Transformers model for DNA-language in genome. Bioinformatics, 37(15):2112–2120, 2021. doi: 10.1093/bioinformatics/btab083.

[6] Jeffry Johnson, Matthijs Douze, and Hervé Jégou. Billion-scale similarity search with GPUs. IEEE Transactions on Big Data, 7(3):535–547, 2019. doi: 10.1109/TBDATA.2019.2921729.

[7] Camille Marchet, Christina Boucher, and Rayan Chikhi. A review of k-mer-based methods for “alignment-free” sequence comparison, 2021.

[8] Brian D. Ondov, Todd J. Treangen, Páll Melsted, Adam B. Mallonee, Nicholas H. Bergman, Sergey Koren, and Adam M. Phillippy. Mash: fast genome and metagenome distance estimation using MinHash. Genome Biology, 17(1):132, 2016. doi: 10.1186/s13059-016-0997-x.

[9] Teresita M. Porter and Mehrdad Hajibabaei. Over 2.5 million DNA barcodes from the International Barcode of Life Project. Scientific Data, 6(1):180308, 2018. doi: 10.1038/sdata.2018.308.

[10] Sujeevan Ratnasingham and Paul D. N. Hebert. BOLD: The Barcode of Life Data System (http://www.barcodinglife.org). Molecular Ecology Notes, 7(3):p355–364, 2007. doi: 10.1111/j.1471-8286.2007.01678.x.

[11] Torbjørn Rognes, Tomáš Flouri, Ben Nichols, Christopher Quince, and Frédéric Mahé. VSEARCH: a versatile open source tool for metagenomics. PeerJ, 4:e2584, 2016. doi: 10.7717/peerj.2584.

[12] Eric W. Sayers, Evan E. Bolton, J. Rodney Brister, Kathi Canese, Joyce Chan, Donald C. Comeau, Robert Connor, Kim Funk, Chaya Kelly, Sunghwan Kim, et al. Database resources of the National Center for Biotechnology Information. Nucleic Acids Research, 50(D1):D20–D26, 2022. doi: 10.1093/nar/gkab1112.

[13] Pierre Taberlet, Eric Coissac, François Pompanon, Christian Brochmann, and Eske Willerslev. Towards next-generation molecular ecology. Molecular Ecology, 21(9):2045–2050, 2012. doi: 10.1111/j.1365-294X.2012.05545.x.

[14] Susana Vinga and Jonas Almeida. Alignment-free sequence comparison–a review. Bioinformatics, 19(4):513–523, 03 2003. ISSN 1367-4803. doi: 10.1093/bioinformatics/btg005. URL https://doi.org/10.1093/bioinformatics/btg005.

